# Gut microbiota of the pine weevil degrades conifer diterpenes and increases insect fitness

**DOI:** 10.1101/116020

**Authors:** Aileen Berasategui, Hassan Salem, Christian Paetz, Maricel Santoro, Jonathan Gershenzon, Martin Kaltenpoth, Axel Schmdit

## Abstract

The pine weevil (*Hylobius abietis*), a major pest of conifer forests throughout Europe, feeds on the bark and cambium, tissues rich in terpenoid resins that are toxic to many insect herbivores. Here we report the ability of the pine weevil gut microbiota to degrade the diterpene acids of Norway spruce. The diterpene acid levels present in ingested bark were substantially reduced on passage through the pine weevil gut. This reduction was significantly less upon antibiotic treatment, and supplementing the diet with gut suspensions from untreated insects restored the ability to degrade diterpenes. In addition, cultured bacteria isolated from pine weevil guts were shown to degrade a Norway spruce diterpene acid. In a metagenomic survey of the insect’s bacterial community, we were able to annotate several genes of a previously described diterpene degradation (*dit*) gene cluster. Antibiotic treatment disrupted the core bacterial community of *H. abietis* guts and eliminated nearly all *dit*-genes concordant with its reduction of diterpene degradation. Pine weevils reared on an artificial diet spiked with diterpenes, but without antibiotics, were found to lay more eggs with a higher hatching rate than weevils raised on diets with antibiotics or without diterpenes. These results suggest that gut symbionts contribute towards host fitness, but not by detoxification of diterpenes, since these compounds do not show toxic effects with or without antibiotics. Rather the ability to thrive in a terpene rich environment appears to allow gut microbes to benefit the weevil in other ways, such as increasing the nutritional properties of their diet.

## Introduction

The interactions between plants and insects are often mediated by plant secondary metabolites. In addition to acting as feeding attractants, these molecules can also be involved in plant defense acting as feeding deterrents or toxins that disrupt gut membranes, impede digestion, hinder normal metabolism, or block ion and nutrient transport, among other effects (Mithoefer and Boland 2012). Herbivores have, in turn, evolved different mechanisms to overcome the noxious effects of plant defenses (Hammer and Bowers 2015). These involve modification of feeding behavior to avoid ingesting high amounts, manipulation of plant defenses to lower their concentrations, increased excretion rates, sequestration away from sensitive processes, target site insensitivity and metabolic degradation (Després et al. 2007).

In addition, insects do not face the threat of plant defenses alone, but together with their gut microbes (Hammer and Bowers 2015). Symbiotic microorganisms have repeatedly been demonstrated to influence interactions between plants and herbivores by supplementing essential nutrients or degrading complex dietary polymers (Douglas 2009). Recently, these microbial functions have been expanded to encompass the manipulation (Chung et al. 2013) and degradation of plant secondary metabolites (Hammerbacher et al. 2013, De Fine Licht et al. 2012 Kohl et al. 2014, Ceja-Navarro et al. 2015, Welte et al. 2015). However, gut microbes are also susceptible to some plant toxins (Bakkali et al. 2008) and might even be their primary target (Mithoefer and Boland 2012).

Many herbivores exploit conifers as a food source. Feeding on conifer tissues is not only problematic nutritionally due to their low concentrations of nitrogen, phosphorus, vitamins and sterols, but conifers typically also contain high concentrations of defensive phenolic compounds and terpenoid resins composed mainly of monoterpene olefins and diterpene resin acids (Keeling and Bohlmann 2006). These defenses are constitutively expressed and can also be induced upon herbivory. With more than 50,000 compounds, terpenes are the most diverse family of plant defenses described to date (Conolly and Hill 1991). Their protective effects include antimicrobial properties (Rastogi et al. 1998, Lunde et al. 2000) as well as feeding deterrence and toxicity against insects, and thus might be involved in conifer resistance against herbivores. Although the exact mode of terpene action is unknown in most cases (Gershenzon and Dudareva 2007), they seem to derive some of their toxic properties from the disruption of gut membranes due to their lipophilic nature or by causing neural damage through compromised ion channels (Keeling and Bohlmann 2006).

Although many insects are able to perform well in the presence of low terpene concentrations, higher amounts can act as deterrents or growth inhibitors (Zhao et al. 2011). A number of terpenes have been associated with toxicity to conifer-feeding insects (Cook and Hain 1988, Raffa and Smalley 1995, Werner 1995). For example, monoterpenes and diterpenes are correlated with white and Sitka spruce resistance against the white pine weevil (*Pissodes strobi*) (Harris et al. 1983, Tomlin et al. 1996, 2000) and diterpenes are correlated with Jack pine resistance against sawflies (Ikeda et al. 1977). Likewise, the Douglas fir pitch moth (*Synanthedon novaroensis*) is more successful attacking lodgepole pines containing low amounts of the monoterpene delta-3-carene (Rocchini et al. 2000). Similarly, the application of methyl jasmonate on seedlings, which increases chemical defenses including terpenes in many conifer species (Martin et al. 2002, Heijari et al. 2005, Schmidt et al. 2005, Moreira et al. 2009) correlates with higher resistance against the pine weevil (Heijari et al. 2005, Erbilgin et al. 2006, Sampedro et al. 2011).

Some environmental bacteria are known to degrade terpenes. For instance, *Pseudomonas abietaniphila* BKME-9 (Martin et al. 2000) and *Burkholderia xenovorans* (Smith et al. 2007) have been reported to degrade diterpene resin acids. However, the ability to degrade and utilize terpenes is not limited to environmental microorganisms, but also occurs in symbiotic microbes associated with vertebrates and invertebrates. Goat rumen-associated bacteria can degrade several monoterpenes (Malecky et al. 2012), and some members of the gut community of bark beetles are capable of *in vitro* degradation of monoterpenes and diterpenes (Boone et al. 2013, Xu et al. 2015). However, whether bacterial degradation of terpenes occurs within the insects as well has not been explored.

The pine weevil, *Hylobius abietis* (Coleptera: Curculionidae: Molytinae), feeds on bark and phloem of several conifer species. It is considered the most damaging conifer pest in managed forests in Europe (Leather et al. 1999, Nordlander et al. 2011) given its devastating impact on newly planted seedlings (Petersson and Orlander 2003). While adults feed both above and below ground, larvae complete their development underground tunneling in the bark of stump roots (Nordlander et al. 2005, Wallertz et al. 2006). Therefore, pine weevils encounter high concentrations of resin terpenes throughout their life cycle. Despite many reports on the noxious effects of terpenoids on herbivores, the pine weevil seems able to cope well with these compounds. Weevils feeding on Sitka spruce show a positive correlation between adult feeding, larval development and high concentrations of both carbohydrates and resin (Langström and Day 2004), suggesting that the pine weevil is adapted to a wide range of terpene concentration. However, high resin content often deters them from feeding in Scots pine (Ericsson et al. 1988). At present, it is not known what mechanisms pine weevils employ to circumvent these compounds, and whether they do so on their own or through association with symbiotic microorganisms.

Here we investigate whether the microbial associates of the pine weevil can play a role in overcoming conifer defenses. The gut microbiota of the pine weevil is geographically stable across Europe, especially within the most abundant bacterial family, the Enterobacteriaceae (Berasategui et al. 2016). The core microbiota of this insect consists of members of the genera *Erwinia, Rahnella,* and *Serratia.* This community assembly seems to be shaped, at least in part, by the nutritional resource these insects exploit (i.e. conifer bark and cambium). It extends beyond the pine weevil, being similar in other conifer feeding beetles, including several species of the genus *Dendroctonus* and *Ips pini* (Cardoza et al. 2006, Adams et al. 2013, Hu et al. 2014, Berasategui et al. 2016, Dohet et al. 2016) as well as wood-feeding wasps (Adams et al. 2011), but it is absent in weevils feeding on non-conifer food sources such as crops or ornamental plants (Berasategui et al. 2016). Among the different members of this conserved community, species of the genera *Pseudomonas*, *Rahnella* and *Serratia* have been described based on metagenomics data (Adams et al. 2013) or phylogenetic inference (Berasategui et al. 2016) to contain many genes involved in diterpene degradation.

In this communication, we follow up on our broad survey of the gut community in the pine weevil and ask whether this consortium of microbes is involved in conferring resistance against plant terpenoid defenses encountered by their insect host. After first finding that diterpene resin acids are reduced during passage through the pine weevil, we manipulated the gut microbiota using antibiotics to determine if microbes are involved in degradation. We also used cultured pine weevil microbes to establish their diterpene degradative capacity outside the insect. Additionally, we sequenced the bacterial metagenome of individuals feeding on different diets in order to explore the taxonomic changes occurring under antibiotic treatment and whether the bacterial community contains the aforementioned diterpene-degrading dit-gene cluster. Finally, we performed bioassays to assess the effect of both diterpenes and gut microbes on the fitness of the insect host.

## Results

### Diterpene levels after passage through weevils

We first determined whether diterpene resin acids were degraded during transit through the pine weevil gut. Insects were fed on Norway spruce branches overnight and their feces were collected after 24 hours. The diterpenes in both food and feces were identified and quantified by gas chromatography-mass spectrometry (GC-MS) of the methyl ester derivatives. Our results show that total diterpene content was reduced by 80% in feces relative to ingested material (Fig. 1A. All individual diterpenes detected were reduced by more than 50%, except dehydroabietic acid (DHAA), which was reduced by only 7% (Fig. 1B). These results demonstrate substantial diterpene degradation during gut passage.

**Figure 1.**
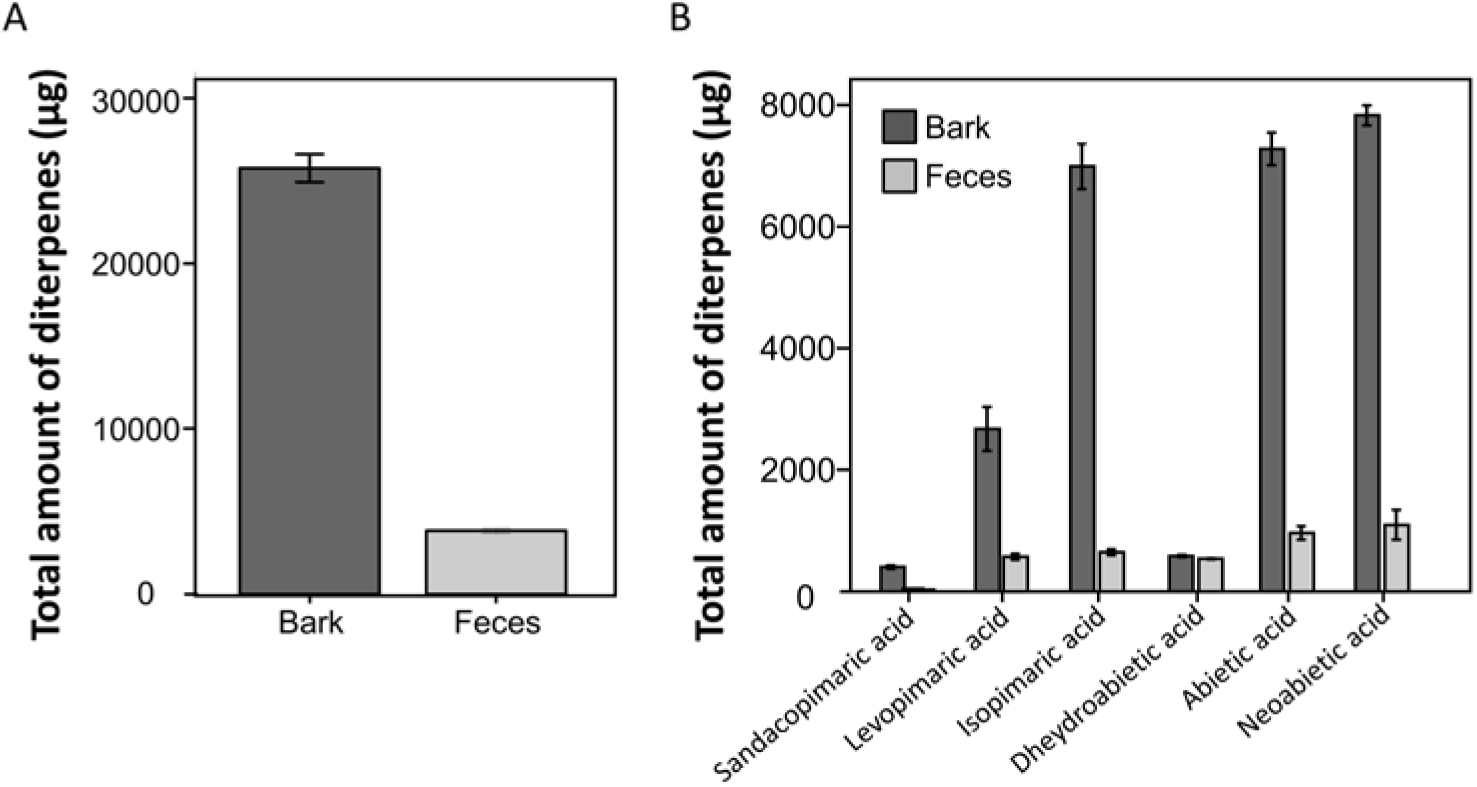
Amounts of (A) total and (B) individual diterpenes ingested by *H. abietis’* feeding on Norway spruce bark (62 insects in 16 hours) compared to the amounts in their feces after digestion. The diterpene content of bark and feces samples was analyzed by GC-MS. The amount of bark ingested was determined by weighing twigs offered to the insect before and after the feeding period. Experiment was repeated twice with similar results.

### Diterpene degradation by bacteria in weevils

To test whether microorganisms mediate weevil degradation of the diterpene resin acids observed after passage through the pine weevil gut, we manipulated the gut microbes through the application of the broad-spectrum antibiotic rifampicin to a semi-artificial diet, and subsequently assessed the diterpene content in the feces of the experimental animals by GC-MS. We observed an increase in the amounts of all the major diterpenes detected in the feces of antibiotic-treated individuals relative to the untreated control group (Fig. 2) (P=0.001 in all cases). Reinfection of the gut with the native community by supplementing the diet with a gut suspension of untreated individuals into the diet of antibiotic-treated insects rescued the insect’s biodegradative capacity, suggesting that gut microbes are responsible for the breakdown of such compounds within the host. This re-acquired degradation capacity resulted in the complete elimination of all diterpenes.

**Figure 2.**
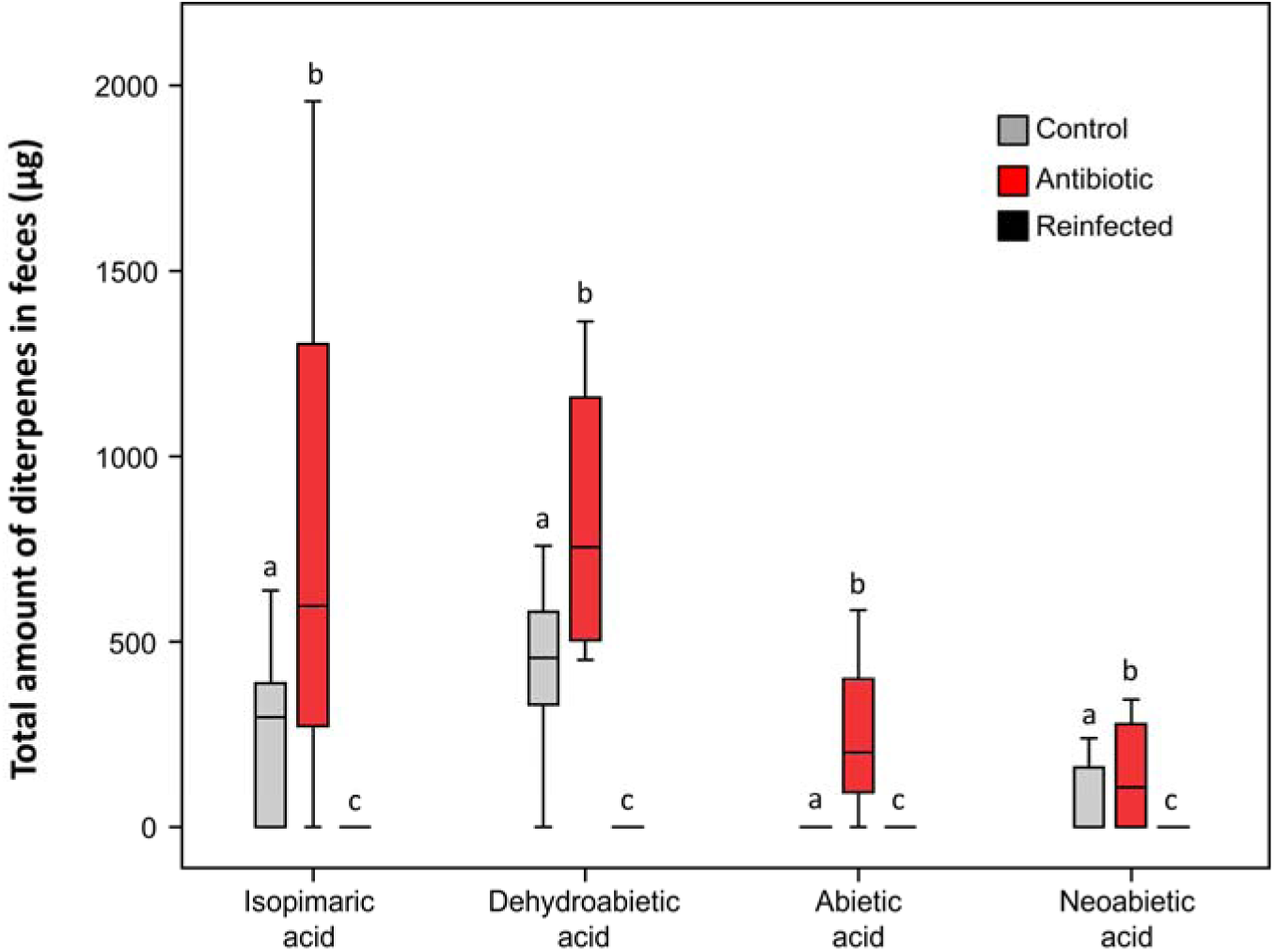
Amount (μg) of major diterpenes in feces of pine weevils feeding on different diets. Color of boxes signifies the experimental treatment: Control, semi-artificial diet with ground Norway spruce for 14 days; Antibiotic, diet amended with rifampicin for 14 days, Reinfected, after being fed with an antibiotic diet for 7 days, these insects were fed diet amended with a weevil gut suspension for 7 days; Lines represent medians, boxes comprise the 25-75th percentiles, and whiskers denote the range.

### Diterpene degradation by cultured weevil microbes

To test whether the gut community of *H. abietis* can degrade terpenes, we isolated bacteria from the guts of the weevil and cultivated them in LB medium overnight. Bacteria were inoculated into medium amended with DHAA (sodium salt) at a concentration of 500 μg/ml, which is in the range of that found in spruce tissue. This compound was chosen because of its commercial availability in high purity and comparatively greater stability in solution, as compared to other diterpene acids from Norway spruce. Sterile, uninoculated medium amended with DHAA served as the control. Although no significant differences in concentration were detected after 1 day of treatment (ANOVA, P=0.98), we observed a significant reduction in the amount of DHAA in the presence of bacteria compared to controls at five days (ANOVA, P=0.03) (Fig. 3).

**Figure 3.**
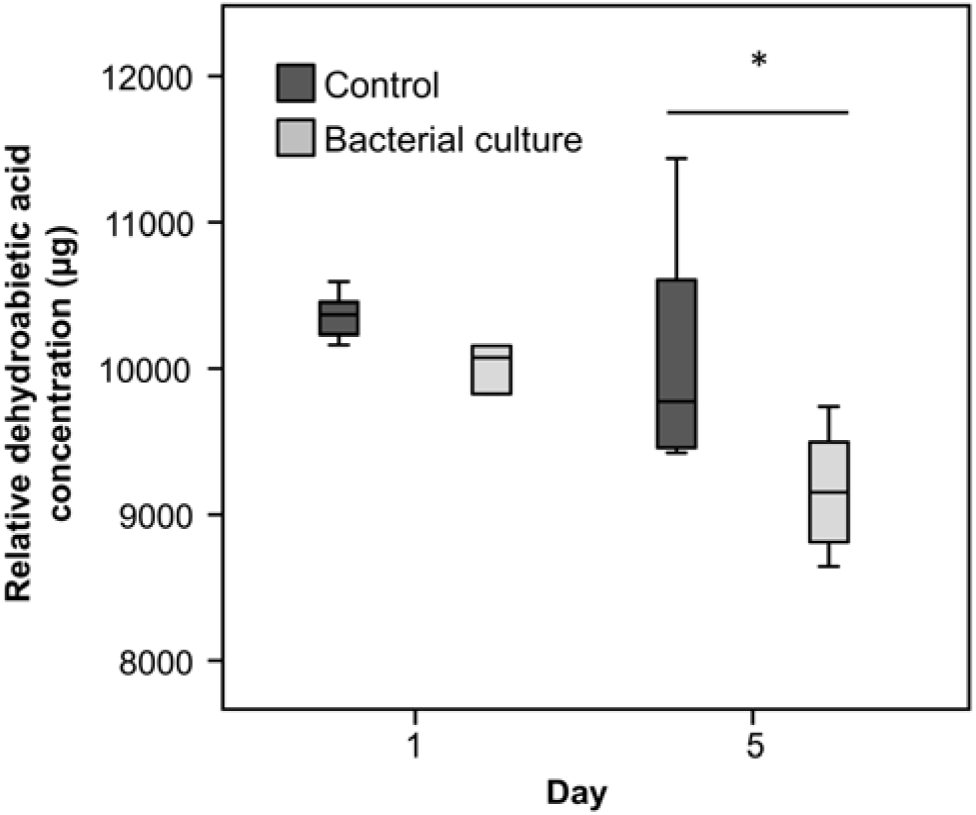
Degradation of the sodium salt of dehydroabietic acid (DHAA) by cultured bacteria isolated from *H. abietis* after 1 and 5 days of growth in LB medium. The initial concentration of DHAA was 14.8μg/mL. Lines represent medians, boxes comprise the 25-75 percentiles, and whiskers denote the range. (ANOVA, P=0.03).

### Metagenomic insights into diterpene degradation

To characterize the genetic basis of bacterial-mediated diterpene degradation in *H. abietis,* we sequenced the bacterial metagenome of weevils feeding on their natural food source (Norway spruce) compared to weevils feeding on an artificial diet, as well as an artificial diet supplemented with antibiotics. Each library contained on average 15.3 million base pairs (Table S1), and assemblies resulted in an average of 49.983 contigs per library.

Of the contigs from the metagenome assembly of weevils fed on spruce, 72.1% were assigned to a bacterial origin, and 27.5% to an eukaryotic origin (Table S2). Consistent with our previous 16S rRNA-based survey of the insect’s microbiome (Berasategui et al. 2016), community profiling using protein-coding genes revealed the gut microbiota of weevils feeding on a coniferous diet to be dominated by gammaproteobacterial associates that could be assigned to various Enterobacteriaceae genera (91.3%; Fig. 4A) including *Erwinia, Rahnella* and *Serratia* (Fig. 4B). All genera that appear in more than 1% abundance belonged to the Enterobacteriaceae family, and nine out of fourteen were previously reported to be present in the pine weevil’s community (Berasategui et al. 2016).

**Figure 4.**
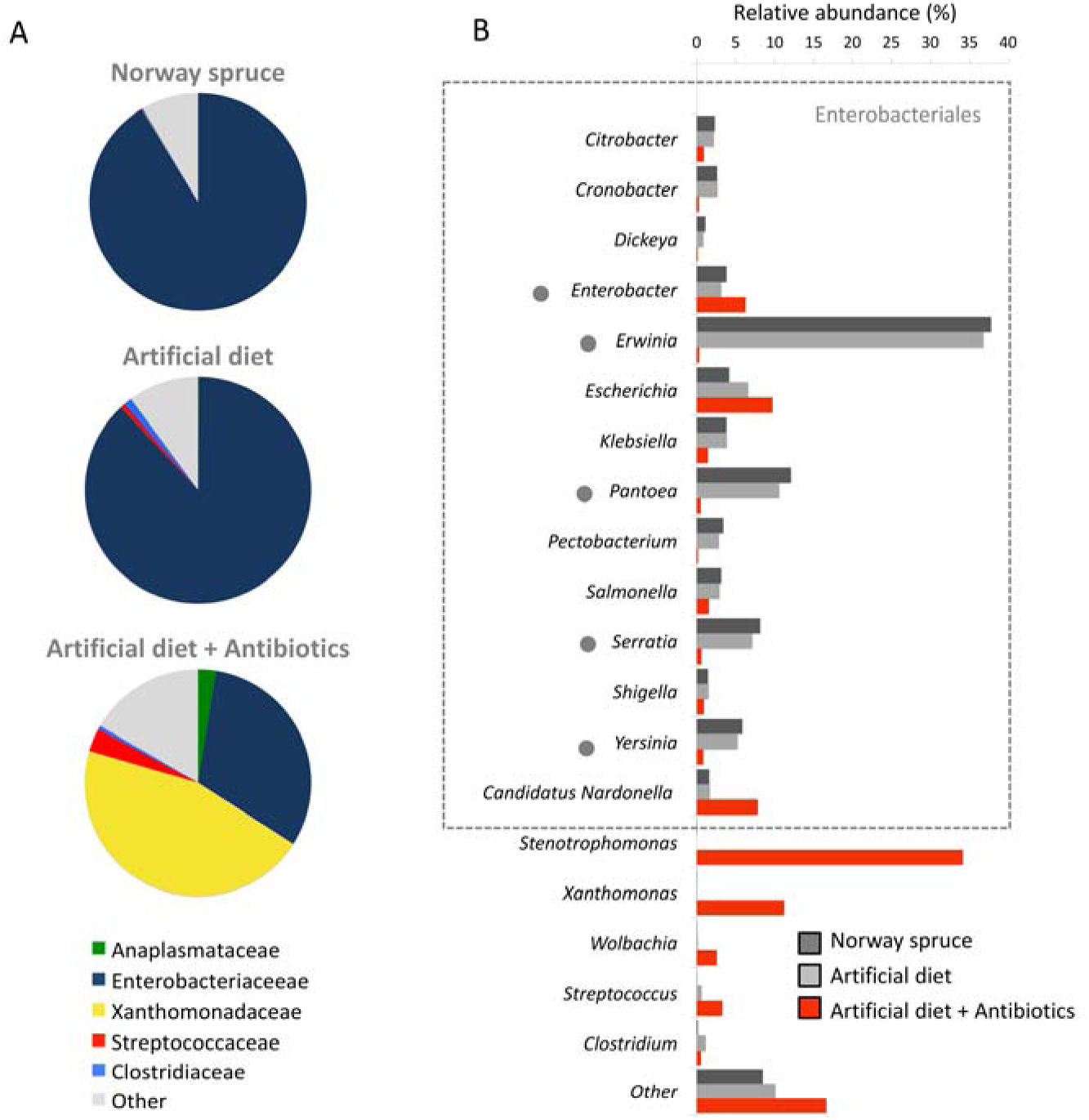
A) Family classification of the bacterial metagenome of pine weevils fed on Norway spruce, artificial diet, and artificial diet with antibiotics, respectively. B) Abundance of different genera in the gut microbiota of weevils feeding on the three diets. Grey dots depict taxa previously found in the gut microbiota of the pine weevil (Berasategui et al. 2016). The dashed box indicates members of the Enterobacteriaceae family.

The gut bacterial community of weevils reared on the artificial diet was qualitatively very similar to that fed on spruce (Fig. 4A). The Enterobacteriaceae was the most abundant family (87.9%) followed by the Streptococcaceae and Clostridiaceae families (1.7% in total) (Fig. 4A). Additionally, all Enterobacteriaceae genera present in more than 1% abundance in spruce-fed weevils were present in artificial diet-fed insects, with only the addition of *Streptococcus.* However, supplementing antibiotics into the artificial diet altered the microbiota, rendering it much more diverse (Fig. 4A), with five families having abundances higher than 1%. In this group, the Xanthomonadaceae family is present in the highest level (45.5%), followed by the Enterobacteriaceae (31.5%), Streptococcaceae (3.29%), Clostridiaceae (0.54%), Anaplasmaceae (specifically *Wolbachia)* (2.5%) and others (16.65%). The Enterobacteriaceae family is the most susceptible to antibiotic treatment, suffering a drastic reduction in overall abundance compared to insects reared on their natural diet or an artificial diet devoid of antibiotics (Fig. 4B).

In order to explore the genetic basis of symbiont-mediated diterpene degradation in the pine weevil, the three bacterial metagenomes sequenced in this study were screened for the presence of a gene cluster (*dit*) predicted to be involved in the degradation of diterpenes in *Pseudomonas abietaniphila* BKME-9 (Adams et al, 2013, Martin and Mohn 2000, Smith et al. 2007, 2008). Genes encoding enzymes that have been previously implicated in diterpene degradation were identified using BLASTn. The metagenome of beetles feeding on Norway spruce contained 10 out of 19 *dit* genes (Fig. 5), and the same was true for beetles reared on the artificial diet, consistent with the minimal changes observed in the composition of the microbial community. However, supplementing antibiotics into the artificial diet led to a near complete loss of *dit-genes* in addition to disrupting the composition of the bacterial associates of *H. abietis* (Fig. 5).

**Figure 5.**
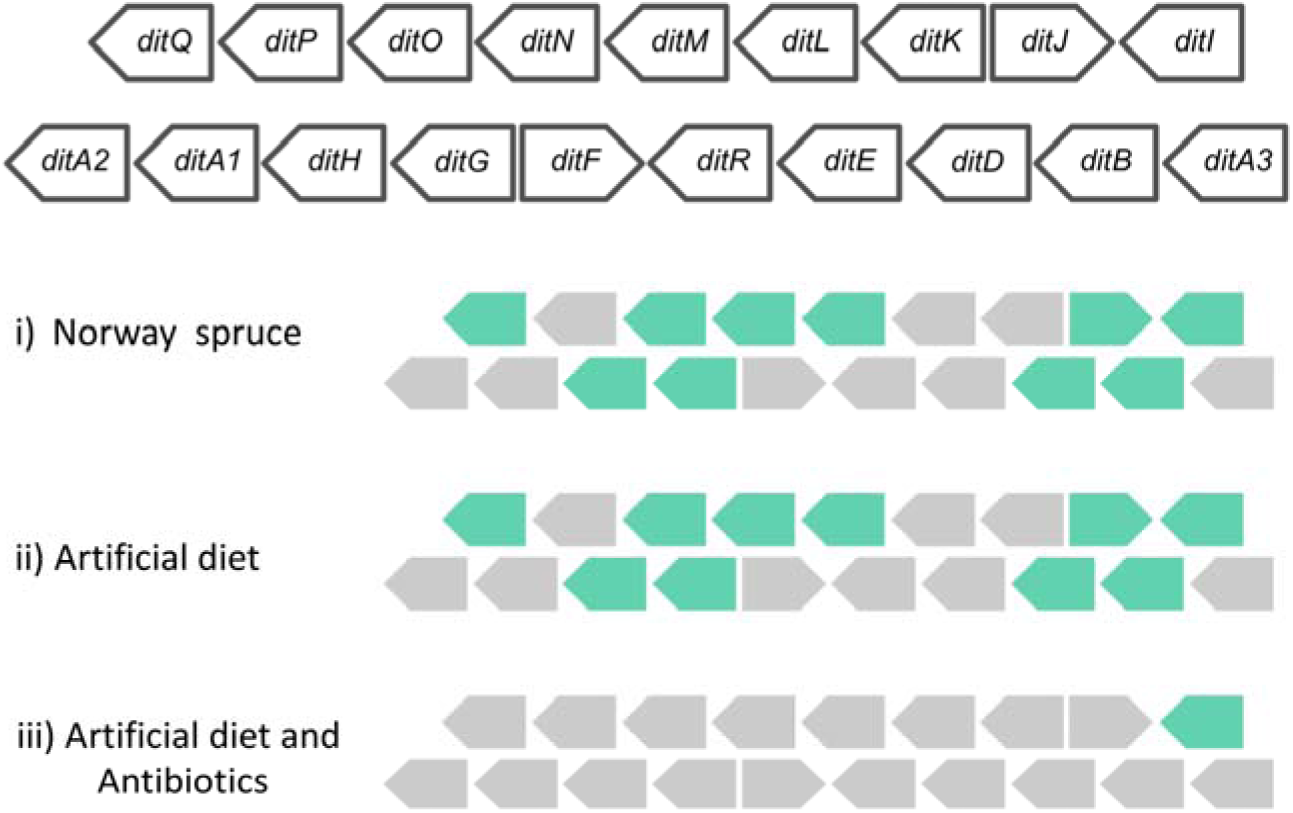
Diterpene gene cluster (Martin and Mohn. 2000) present in the metagenome of pine weevils feeding on different diets: Norway spruce, artificial diet, and artificial diet amended with antibiotics. Coloring: turquoise, present; grey, absent.

### Effect of diterpenes and gut bacteria on weevil fitness

In order to assess the effect that diterpenes and microbially-mediated diterpene degradation might have on pine weevil fitness, we measured survival and fecundity of weevils feeding on different types of artificial diet. Insects were fed on diet with a natural mixture of diterpene acids, with the antibiotic rifampicin, with both diterpenes and antibiotic, and without any supplementation.

We observed no difference in survival rates depending on treatment (Mantel-Cox P=0.18; Breslow P=0.27; Tarone-Ware P=0.19) or sex (Mantel-Cox P=0.56; Breslow P=0.67; Tarone-Ware P=0.66) after 30 days (Fig. 6A). However, we observed differences between treatments in relation to the number of eggs laid (Fig. 6B). Individuals that fed on diet with diterpenes only (no antibiotic added) laid more eggs than individuals in any of the other three groups (P=0.05). Likewise, hatching rates of eggs laid by diterpene-fed mothers with their native microbiota were significantly higher than in any of the three other groups (P=0.01, Fig. 6C).

**Figure 6.**
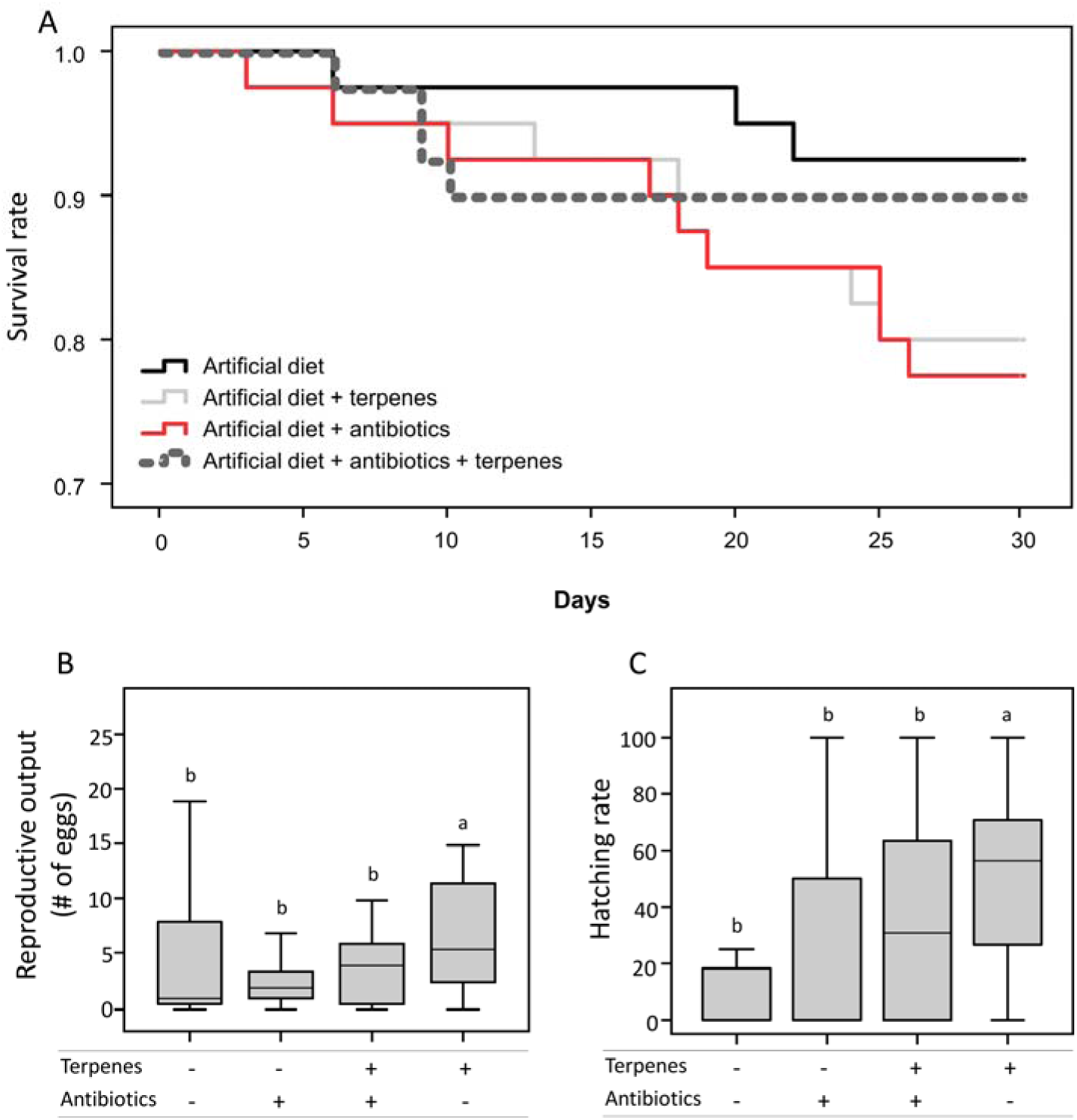
Fitness parameters of pine weevils fed on artificial diet containing no additives (black), 0.3% (w/v) of the antibiotic rifampicin (red), 3% (w/w) of a natural diterpene mixture (grey), and the antibiotic plus diterpenes (dashed). In the box plots, lines represent medians, boxes comprise the 25-75 percentiles, and whiskers denote the range. (a) Survival: treatment (Mantel-Cox P=0.18; Breslow P=0.27; Tarone-Ware P=0.19); sex (Mantel-Cox P=0.56; Breslow P=0.67; Tarone-Ware P=0.66). (b) Number of eggs laid (ANOVA, P=0.05) and (c) egg hatching rate (ANOVA, P=0.01).

## Discussion

Microorganisms associated with herbivorous insects have been shown to mediate the outcome of insect-plant interactions across a number of insect groups in different ways (Tsuchida et al. 2004, Hosokawa et al. 2007). For example, microorganisms can aid in the exploitation of plant resources by supplementation of essential nutrients, degradation of complex structural metabolites (Douglas 2009) and manipulating (Chung et al. 2013) or overcoming (Hammer and Bowers 2015) plant defenses. Microbial symbionts of herbivorous animals have long been suspected to aid in the detoxification of plant secondary metabolites and thereby contribute to host fitness, but direct experimental evidence remains scarce (but see Kohl et al. 2014, Ceja-Navarro et al. 2015). Here we show that diterpene resin acids, a group of plant defense compounds abundant in the bark of conifer trees, are degraded in the pine weevil after ingestion. Gut microbes of the pine weevil contain genes encoding enzymes that catalyze the degradation of diterpene acids. These microbes were found to degrade diterpenes in the insect and when cultured separately on medium containing natural concentrations of one diterpene acid. Diterpene supplementation of the pine weevil diet in the presence of the native gut microbiota enhances the weevil’s fecundity and egg hatching rate.

### Diterpene-degrading genes in the pine weevil gut bacterial metagenome nearly eliminated by antibiotic treatment

In agreement with our previous report using 16S rRNA profiling (Berasategui et al. 2016), taxonomic classification of the gut bacterial metagenome of the pine weevil revealed a community dominated by the Enterobacteriaceae family. Specifically, members of *Erwinia, Pantoea, Serratia,* and *Yersinia,* encompassed 90% of the whole microbiota. Although we detected *Wolbachia* species in all three metagenomes sequenced, their abundance was markedly lower than that previously reported using 16S rRNA data (Berasategui et al. 2016). While differences in *Wolbachia* titers within a single insect population have been previously described (Müller et al. 2013), it is possible that variation in this case arises from bias in the PCR reaction required for 16S pyrosequencing, which is not required for metagenome sequencing. The very low abundance of Firmicutes in the present study is consistent with that of weevils previously collected in the same location (Spain) which were devoid of this bacterial family (Berasategui et al. 2016). However, members of the Enterobacteriaceae were as dominant as in the previous study supporting their role as stable and conserved constituents of the pine weevil gut microbiota in contrast to the variability affecting other members of the community. The conserved fraction of the pine weevil microbiota overlaps with that of other conifer-feeding beetles as well as a conifer-feeding wasp (Adams et al. 2011) suggesting that this is shaped by a shared environment and may be adaptively significant (Berasategui et al. 2016).

The pine weevil microbial community remained unaltered after two weeks of beetles feeding on an artificial diet (Fig. 4A), indicating that it is also resilient to changes in diet. However, the addition of antibiotics did alter community composition, reducing the relative abundance of most Enterobacteriales (with the exception of *Enterobacter* sp., *Escherichia* sp., and the bacteriome-localized endosymbionts), and increasing that of *Stenotrophomonas* sp., *Xanthomonas* sp., and *Wolbachia* (Fig. 4B).

Our previous inference of metagenomic functions via PICRUSt (Berasategui et al. 2016) suggested the enrichment of a gene cluster known as *dit* among *H. abietis’* bacterial associates that is potentially involved in diterpene degradation based on studies of free-living bacteria (Adams et al. 2013, Martin et al. 2000, Smith et al. 2007, 2008). Functional annotation of the bacterial metagenome in the present study revealed the presence of ten out of the 19 known *dit* genes in weevils feeding on their natural food source as well as on an artificial diet (Fig. 5). However, the supplementation of antibiotics in the artificial diet resulted in the loss of all but one *dit* gene (Fig. 5).

Even with only 10 of 19 reported *dit* genes, the members of the *H. abietis* microbial community may still be able to degrade diterpene resin acids. Not all 19 genes are required for bacterial catabolism of diterpenoids by *Pseudomonas abietaniphila* BKME-9 (Martin and Mohn 2000, Smith et al. 2004, 2007). For example, knocking out *ditR* does not impair the growth of this *Pseudomonas* strain on diterpene-rich media (Martin and Mohn 2000). Conversely, knocking out *ditQ,* restricts growth of *P. abietaniphila* on dehydroabietic acid but not on abietic acid (Smith et al. 2004). Moreover, *ditI* and *ditH,* both reported as essential genes for diterpene degradation, were found to be present in our metagenomic survey of the *H. abietis’* gut bacterial community (Fig. 5). Parallel to previous findings (Adams et al. 2013), phylogenetic binning of dit-gene sequences annotated within our metagenomes revealed that most of the sequences belonged to taxa from the Enterobacteriaceae, strongly supporting the involvement of this bacterial family in the degradation of diterpenes within *H. abietis’* gut. Despite also belonging to the Enterobacteriaceae, the primary endosymbiont of pine weevils, *Nardonella* sp. (Conord et al. 2008), is not likely to play a major role in this degradation given that it is not affected by antibiotic treatment (Fig. 4B), and thus is not responsible for either the differential occurrence of *dit* genes or the reduction in terpene concentration observed in this study.

### Pine weevil gut microbes degrade diterpenes in the insect and in isolated cultures

The possibility of microbial degradation of diterpenes in the pine weevil was first suggested by our measurements showing an 83% decrease in total diterpene content between the spruce bark ingested by the weevil and the frass. To determine if the microbiota degrades diterpenes inside the insect, we manipulated the gut community through the addition of antibiotics to the insect diet. Our results indicated that the degradation of diterpenes decreased upon addition of antibiotics (Fig. 2), and these capabilities were rescued in antibiotic-treated insects upon supplementing their native bacterial community through the diet. We then tested the ability of isolated weevil microbes to degrade dehydroabietic acid (DHAA) in solution at concentrations found in spruce bark, and found a significant decline (Fig. 3). Thus experiments with both weevils and isolated bacterial illustrate the contribution of gut bacteria towards diterpene breakdown. The correlation of reduced breakdown rates with a reduction in the expression of diterpene catabolizing genes under antibiotic treatment is another line of evidence linking the pine weevil gut bacterial community to diterpene degradation.

Symbiotic bacteria have been previously demonstrated to breakdown conifer resin terpenes. For example, close relatives of *Serratia, Rahnella* (both Enterobacteriales), *Pseudomonas,* and *Brevundimonas* isolated from the gut of bark beetles (*Dendroctonus ponderosae* and *D. valens*) were found to degrade monoterpenes and diterpenes (Boone et al. 2013, Xu et al. 2015). Free-living microbes isolated from conifer pulp mill wastewater and forest soil, such as *Pseudomonas abietaniphila* BKME-9 (from which the *dit* gene cluster was reported) and *Burkholderia xenovorans* LB400, are also described to degrade diterpenes (Smith et al. 2008). These bacteria are also able to utilize diterpenes as their sole carbon source (Martin and Mohn 2000, Morgan and Wyndhan 2002, Smith et al. 2007).

The degradation of plant secondary metabolites by symbiotic bacteria is not limited to terpenes, nor to insects. For instance, members of the gut community of the cabbage root fly *(Delia radicum)* harbor a plasmid (saxA) involved in metabolizing the isothiocyanates of the host plant (Welte et al. 2016). The coffee bean borer (*Hypotenemus hampei*) relies on at least *Pseudomonas fulva,* a member of its gut bacterial community, to degrade the alkaloid caffeine, thereby increasing the insect’s fitness (Ceja-Navarro et al. 2015). Likewise, desert woodrats (*Neotoma lepida*) harbor a gut microbial assembly that allow their hosts to exploit the toxic creosote bush (*Larrea tridentata*) through the degradation of phenolic compounds (Kohl et al. 2014, 2016). In addition to bacteria, symbiotic fungi can also degrade plant secondary metabolites for their hosts. *Lasioderma serricorne,* the cigarette beetle harbors a symbiotic yeast (*Symbiotaphrina kochii*) in its digestive system that is able to degrade several plant toxins and use them as sole carbon sources (Dowd and Shen 1990, Shen and Dowd 1991). Similarly, the phenolics that leaf cutter ants (*Acromyrmex echinatior*) encounter in leaves, are detoxified by a laccase produced by the ant fungal cultivar, *Leucocoprinus gongylophorus* (De Fine Licht et al. 2012).

### Gut bacteria increase pine weevil fitness, but not by direct detoxification of diterpenes

To assess the impact of gut bacteria on the pine weevil performance, we measured several fitness parameters on weevils feeding on diets with and without the diterpenes and with and without the broad-spectrum antibiotic rifampicin. We observed a fitness benefit for the weevil as evidenced by higher fecundity and hatching rate only when insects fed on diterpene-containing diet with a full spectrum of bacteria (without antibiotics) (Fig. 6B and 6C). Different hypotheses might explain how gut bacteria can enhance host fitness through the metabolism of terpenes. For example, removal of diterpenes could reduce the levels of compounds toxic to the weevils, but our bioassays show that insects that lack their full microbial complement do not suffer higher adult mortality when feeding on diterpene-containing diet than insects with all of their native microbes, suggesting that insects have an intrinsic mechanism of circumventing diterpenoid toxins. Microbes could conceivably assist insects in degrading terpene toxins (Raffa 2014), but such a contribution does not seem important in the case of *H. abietis.*

Another explanation for the fitness benefits of microbes is that they could improve the nutritional properties of the pine weevil diet allowing females to allocate more resources to egg production (Wainhouse et al. 2001). Degraded terpenes might be directly employed as substrates for insect respiration. Or, if terpenes are readily used as respiratory substrates by microbes (DiGuistini et al. 2011, Wang et al. 2014), they could enhance microbial growth and so increase the breakdown of refractory carbohydrate polymers or increase supplies of nitrogenous compounds, vitamins or sterols (Douglas 2009, McCutcheon and Moran 2007, Salem et al. 2014), all scarce resources in conifer bark and phloem. For example, Morales-Jiménez and colleagues (2009, 2012) demonstrated that members of the gut community of the conifer-feeding bark beetles *D. ponderosae,* (*Rahnella sp. Pantoea sp* and *Stenotrophomonas sp.*) and *D. rhyzophagous* (*Pseudomonas sp, Rahnella sp.* and *Klebsiella sp.*) can fix nitrogen (Morales-Jimenez et al. 2012) while free-living fungal associates of bark beetles can supply *Dendroctonus* beetles with sterols (Bentz and Six, 2006).

Hence for *H. abietis* the fitness benefits of its gut microbes may not be derived directly from terpene degradation. Rather, the ability to degrade terpenes may allow microbes to thrive in a terpene-rich environment and enhance pine weevil growth by providing critical nutrients and vitamins. Given that the taxonomical composition and functional capabilities of the weevil microbial assembly strongly resemble those of other bark beetles, wood-feeding wasps, and sawflies exploiting very similar ecological niches (Berasategui *et al.* 2016, Adams *et al.* 2011, 2013, Boone et al. 2013, Whittome *et al.* 2007), these other insect groups may also accrue such fitness benefits from their gut microbiota.

## Material and methods

### Insect collection and maintenance

Adult pine weevils (*Hylobius abietis*) were collected near Neustadt, Lower Saxony (Germany) and in Galicia (Spain). In Germany weevils were collected by leaving a recently cut log near a clear cut for some days and manually collecting the insects that were attracted. In Spain beetles were collected with clean pit-fall traps baited with α-pinene and ethanol (Nordlander 1987). Vials without lids were filled with ethanol, and the entrance blocked with bait-impregnated paper towels or cotton and placed leaning towards a pine branch. Once in the lab, the beetles were stored in darkness at 10°C in boxes of 50 individuals with moist paper, a container with soil and Norway spruce (*Picea abies*) twigs. Insects were brought to the lab bench one week before each experiment for acclimatization.

### Semi-artificial and artificial diet preparation

#### Semi-artificial diet

Bark and cambium of Norway spruce branches were manually removed from the wood and frozen in liquid nitrogen. Needles were removed with a scalpel and the remaining tissues were homogenized manually with a mortar and pestle under liquid nitrogen. Agar (1.60 g, Roth, Karlsruhe, Germany) was diluted in 50 mL of water and let cool to ~ 60°C. This mixture was added to 18.75 g of the homogenized spruce tissue and stirred until homogeneous. The antibiotic containing semi-artificial diet was prepared by adding the broad spectrum antibiotic rifampicin to the diet to a final concentration of 0.3% (w/v).

#### Complete artificial diet

The diet used for seed-feeding pyrrhocorid bugs described in Salem et al. (2014) was modified with the addition 6 mL of sunflower seed oil instead of 20 mL.

### Degradation of diterpenes on passage through weevils

Pine weevils (62 individuals) were kept without food in plastic boxes for 4 days and sprayed with water daily to maintain humidity. After this starvation period, insects were allowed to feed on Norway spruce twigs *ad libitum.* After 16 hours, insects were transferred to glass petri dishes and allowed to defecate for 24 hours. Feces were collected and suspended in 5 ml water. We added the suspended feces to tert-butyl methyl ether (1:5, v/v) in a total of 25 ml and let the mixture shake for 20 hours, after which 5 ml of 0.1 mM (NH_4_)_2_CO_3_, pH 8.0, were added to the mixture and vortexed (Schmidt et al. 2011). The ether layer was removed and concentrated to 0.5 ml and analyzed for diterpenes by GC-MS (see below) alongside the ether fraction of the bark of Norway spruce twigs prepared in a similar manner. The quantity of bark ingested by the weevils in the feeding period was determined by weighing the twigs before and after with a correction for water loss.

### Manipulation of the gut microbiota and *in vivo* degradation of diterpenes

To assess the potential role of the gut bacteria in the degradation of terpenoids within the beetle gut, six weevils (three males and three females) in individual petri dishes were allocated to each of three treatments: (i) control, (ii) antibiotic, and (iii) reinfected. Control individuals were fed on control semi-artificial diet for 14 days, whereas antibiotic-treated individuals were fed on semi-artificial diet amended with the antibiotic rifampicin for 14 days. Reinfected individuals were fed on the antibiotic diet for half of the experiment and then switched to a semi-artificial diet amended with a gut suspension of untreated weevils for the remaining time. We generated this suspension by crushing the guts of four untreated weevils in 1 ml PBS and vortexing. Insects were provided with 100 mg of diet every day that was supplemented with 10 μL of gut suspension in the case of the reinfected treatment or PBS in the control. Feces were collected daily and frozen at -20°C. Feces from the last experimental day were extracted with 200 μl *tert*-butyl methyl ether, shaken overnight and prepared for diterpene analysis via GC-MS as described below.

### Degradation of diterpenes by cultured gut microbes

Six beetles were dissected under sterile conditions. Individual guts were suspended in a vial of 1 mL of PBS (phosphate-buffered saline). To separate bacteria from the gut walls, tissues were sonicated (50/60 Hz, 117 V, 1.0 Amp) for 30 s, macerated with a pestle and vortexed at medium speed for 10 s. Vials were centrifuged at 3000*g* to pellet host tissues and the supernatant was filtered using 10 μm syringe filters.

A 10 μL quantity of each bacterial suspension was inoculated in 10 mL LB medium and grown overnight shaking at 220 rpm at room temperature. Overnight cultures were diluted to an optical density of 0.1 at 600 nm (OD_600_) with additional LB medium. To test whether the gut community of the pine weevil is able to degrade diterpenes, we inoculated 10 μL of the diluted bacterial culture in a vial of 990 μL LB medium amended with dehydroabietic acid (DHAA) as the sodium salt. In the same way, 10 μL LB medium was inoculated in control vials of medium amended with DHAA. The experiment was carried out in a 24-well flat bottom plate (Sarstedt, Numbrecht Germany), replicated six times, and allowed to grow for 5 days. Samples were taken for analysis on the 1^st^ and 5^th^ days. Upon sampling, the content of each well was transferred to Eppendorf tubes and centrifuged at 3000*g* for 5 min to pellet bacterial cells. The supernatant was then transferred to glass vials for diterpene analysis by HPLC rather than GC-MS to detect the sodium salts directly.

The sodium salt of DHAA was produced by dissolving dehydroabietic acid (Sigma, 1.33 g) in 12 mL MeOH. This mixture was amended with an equimolar amount of NaHCO_3_ (372 mg) dissolved in 3 mL of water. The mixture was left standing in a closed vial for 14 days at room temperature to allow the reaction to proceed. Subsequently, the solution was filtered with a syringe filter and evaporated to dryness under nitrogen. In order to prepare the experimental medium, 750 mg of DHAA in this form was added to 1.5 L of LB medium. After autoclaving (121°C, 20 min), the solution was vacuum-filtered to remove un-dissolved particles.

### DNA extraction and Illumina-based metagenome sequencing

For one week, a group of six weevils were reared on one of three different diets: (1) Norway spruce (*Picea abies*) twigs, (2) artificial diet (AD), or (3) artificial diet amended with the antibiotic rifampicin at 0.3% (w/v). Through dissection, the midgut region was harvested from every individual using sterile forceps and iris scissors. Once dissected, DNA was extracted from the midgut using the Microbiome Kit (Qiagen, Hilden, Germany) following the manufacturer’s instructions. Equimolar concentrations of samples from the same treatment were pooled, resulting in a single sample per treatment. Sequencing of a 150-bp library was conducted for each treatment using an Illumina Genome Analyzer IIx at the Genome Center of the Max Planck Institute in Cologne (Germany). The assembly was generated with Meta-Velvet v1.0.19 (Namiki et al. 2012) based on ~40 million quality-filtered read pairs and subjected to a gap-closing analysis. For annotation of gene content as well as taxonomic assignment, we used the Meta Genome Rapid Annotation using Subsystem Technology (MG-RAST) (Meyer et al. 2008).

We determined the community profile in the metagenomic dataset using the “Best Hit Classification” tool of MG-RAST to classify CDSs on the basis of best BLASTP hits with a minimum identity cutoff of 60%, and a maximum e-value cutoff of 10^−5^. Sequences were classified at the genus level (95% identity), and taxa with <1% abundance were removed. To examine the enrichment patterns of genes encoding diterpene degradation genes across the different treatments, we performed BLASTP against a custom data set of proteins belonging to the *dit* diterpene acid degradation pathway of *Pseudomonas abietaniphila* BKME-9, as previously described by Adams et al. (2013).

### Fitness assays

Weevils (80 male and 80 female) were randomly distributed as pairs in plastic boxes. Each box contained one piece of moisture paper and an Eppendorf tube lid as container for artificial diet prepared as described above. Each box was randomly assigned to one of four treatments: (i) diet without additions, (ii) diet with the antibiotic rifampicin at 0.3% (w/v), (iii) diet with a natural mixture of diterpene acids and (iv) diet with both antibiotic and diterpene acids. The diterpene acids were from a natural conifer mixture, sold as “gum rosin” (Sigma), which was dissolved in methanol, added to the diet as 3% (w/w), and the methanol allowed to fully evaporate before using. Weevils were subjected to the treatments for one month. As fitness parameters, survival, number of eggs laid and number of eggs hatched were recorded every 24 hours.

### Statistics

#### *In vitro* assays

Differences in the concentration of terpenes in liquid cultures were assessed in SPSS with an ANOVA.

#### *In vivo* assays

Differences in the amount of terpenes in the feces of weevils were analyzed using a Kruskal-Wallis test.

#### Fitness assays

The number of eggs laid and the hatching rate were analyzed in R with a generalized linear model. Survival was analyzed in SPSS using three different statistical tests: Mantel-Cox, Breslow, and Tarone-Ware.

### Chemical analyses

#### GC-MS analysis of diterpene resin acids

Diterpene resin acids of Norway spruce bark and pine weevil feces were analyzed as described in Schmidt et al. (2011). In brief, *tert*-butyl methyl ether extracts that had been washed with (NH_4_)_2_CO_3_ to remove small organic acids were methylated with N-trimethylsulfonium hydroxide in methanol and concentrated under nitrogen. Separation was accomplished on a polyethylene glycol stationary phase by split injection. Compounds were identified by comparison with methyl esters produced from authentic diterpene acid standards and quantified with an internal standard of methylated dichlorodehydroabietic acid.

#### HPLC analysis of diterpene resin acids (sodium salts)

Samples were analyzed on an HPLC (1100 series equipment, Agilent Technologies, Santa Clara, CA, USA), coupled to a photodiode array detector (Agilent Technologies). Separation was accomplished on a Nucleodur Sphinx RP column (250 × 4.6 mm, 5 μm particle size, Macherey-Nagel, Düren, Germany). The mobile phase consisted of 0.2% formic acid in water (A) and acetonitrile (B) with a flow of 1 ml per minute with the following gradient: start 20% B, 0-20 min 20-100% B, 20-23 min 100% B, 23.1-28 min 20% B. For diterpene quantification, peaks were integrated at 200 nm and an external standard curve with an authentic standard of DHAA (Sigma, Taufkirchen, Germany) was created.

## Conflict of interests

The authors declare no conflict of interests.

## Author Contributions

AB carried out the microbiological and chemical lab work, performed the data and statistical analyses, and wrote the manuscript; HS carried out the molecular work and metagenome analysis; CP synthetized the DHAA salt; VMS aided in chemical lab work; AS provided reagents; AB, AS, MK, JG and conceived of the study. All authors gave critical comments on the manuscript and gave final approval for publication.

## Acknowledgements

We would like to thank Selina Secinti and Marcus Grant for laboratory assistance. We are grateful to Luis Sampedro, from the Misión Biológica de Galicia (MBG-CSIC) for providing weevils. We gratefully acknowledge funding from the Max Planck Society (all authors), and the Alexander von Humboldt Foundation (HS).

## Data Accesibility

Accession numbers will be provided upon acceptance.

